## Supplemental Figures for "Indisulam synergizes with palbociclib to induce senescence through inhibition of CDK2 kinase activity"

### Supplemental Figure 1

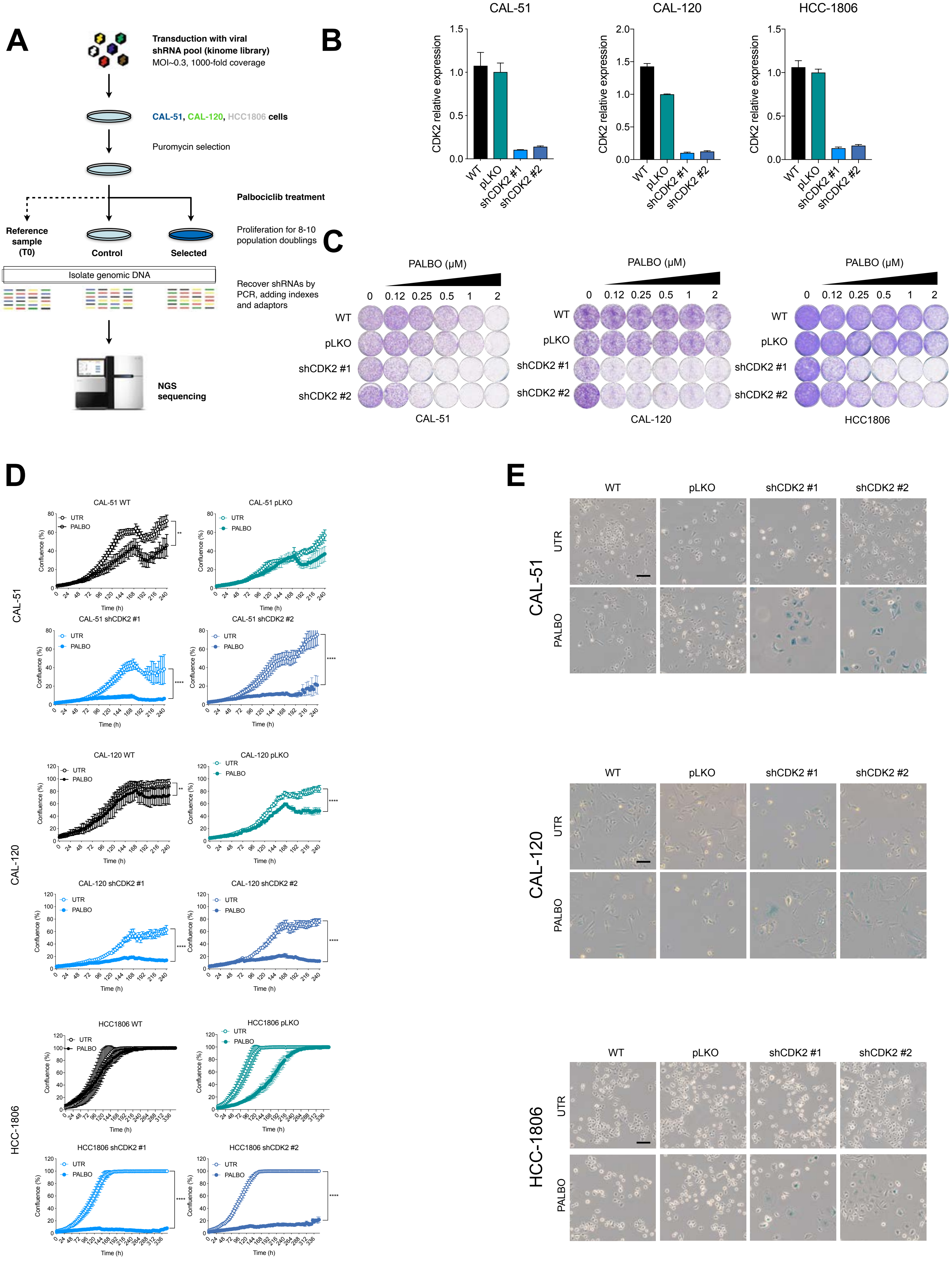

Supplemental Figure2

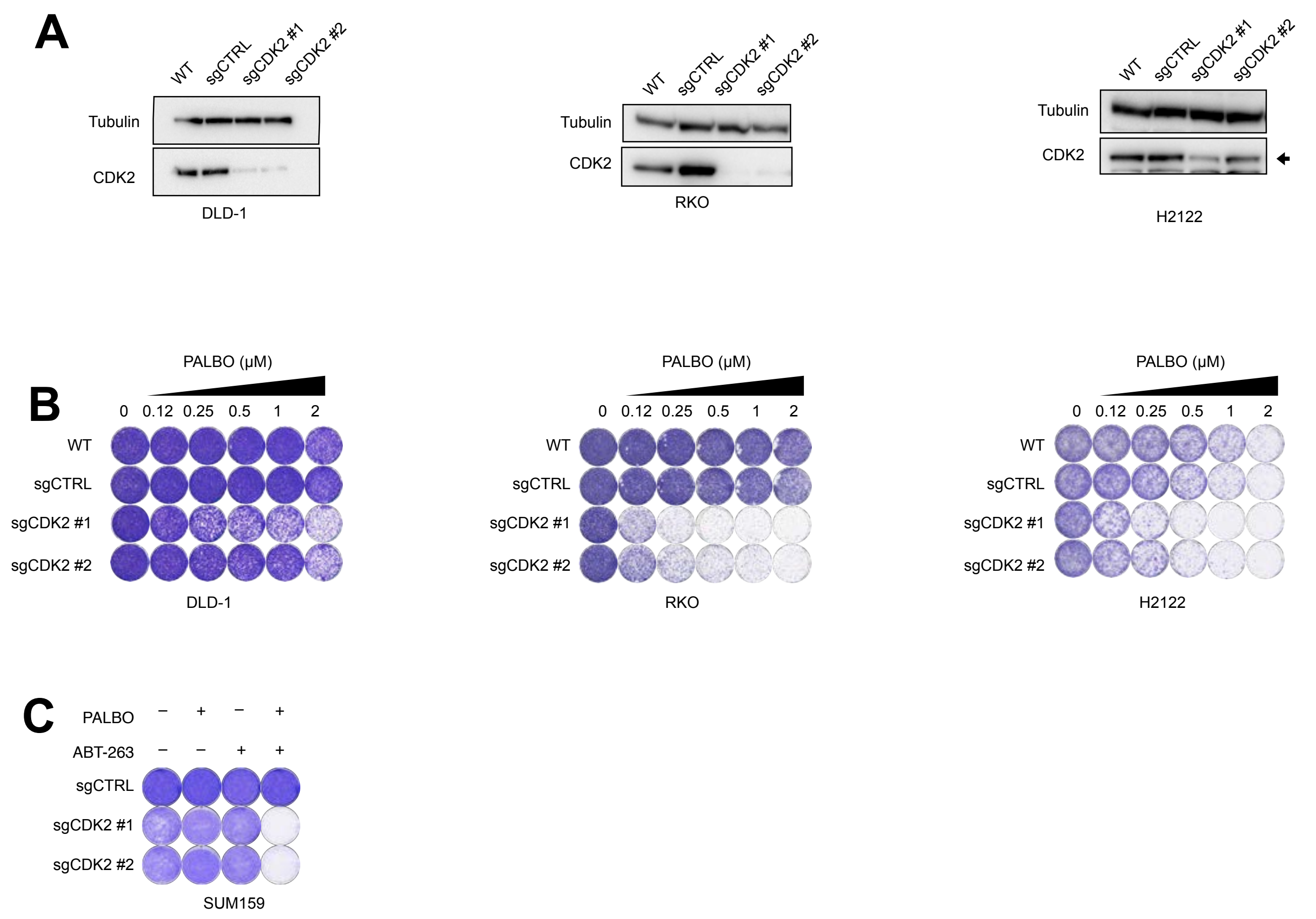

### Supplemental Figure 3

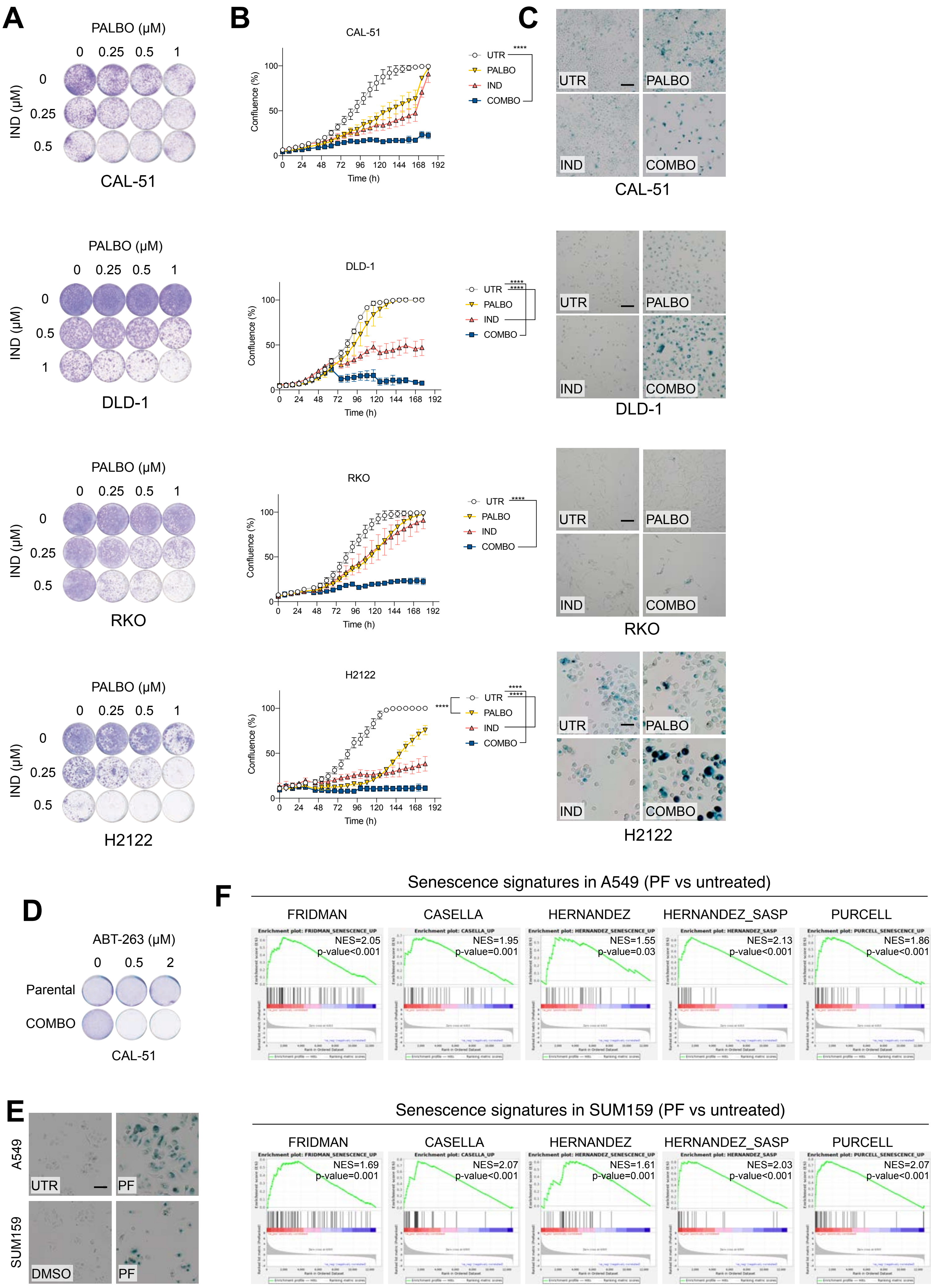

### Supplemental Figure 4

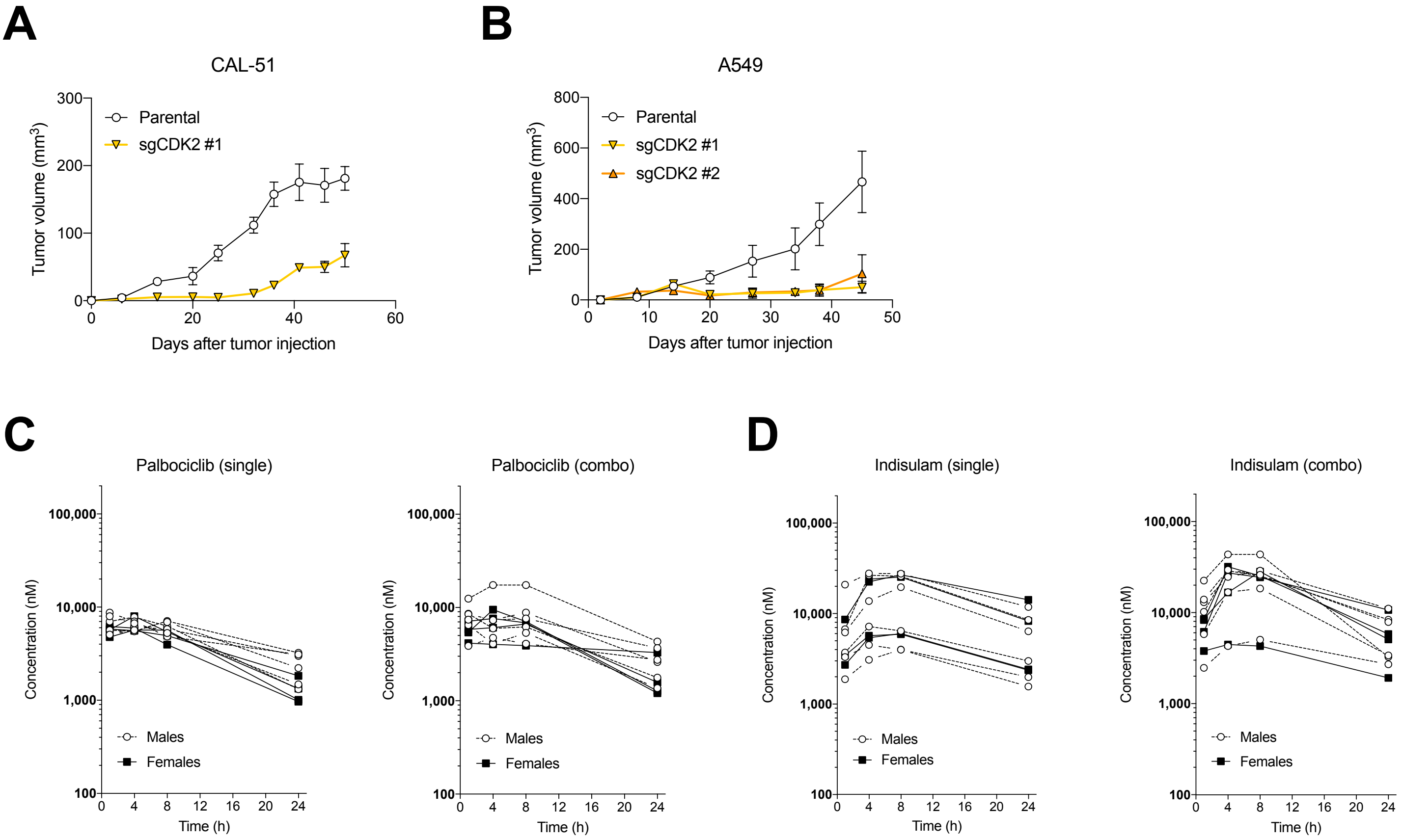

Supplement Figure 5

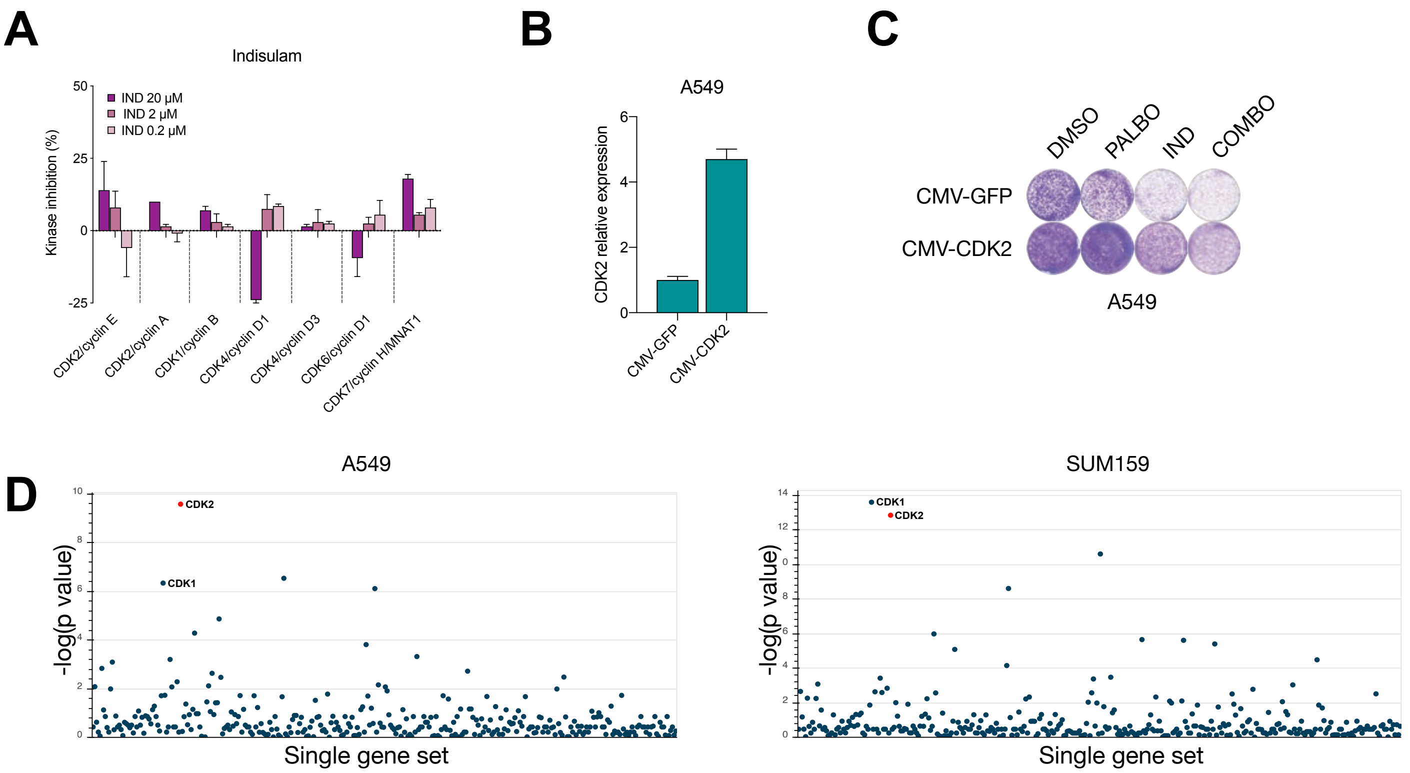
